# Filling the gap in the Neotropics: new firefly species provide insights into Central American biodiversity

**DOI:** 10.1101/2025.02.20.639311

**Authors:** Ana Catalán, Claudia María Pérez Archila, Adriana María Echeverría Méndez, José Andrés Guitiérrez, José Monzón, Viridiana Vega Badillo, Sebastian Höhna

## Abstract

Tropical ecosystems, particularly in the Neotropics, remain vastly understudied, especially regarding the biodiversity of many taxa such as fireflies. This study investigates the biodiversity and phylogenetic relationships of fireflies in Guatemala, focusing on the genera *Photinus, Photuris*, and *Bicellonycha*. Using genetic barcoding we generate time-calibrated phylogenetic trees, revealing deep divergent times between Guatemalan and North American fireflies and potentially multiple colonialization events. Using morphometric information, taxonomic keys and species delimitations models (*mPTP*) we identified four new *Photinus* species: *P. hunahpú, P. helodermensis, P. schusteri*, and *P. semetabajense*. This work highlights the significant firefly biodiversity in Nuclear Central America and the potential for future species discovery. Biodiversity research is necessary for conservation efforts of key ecosystems and will broaden our understanding of general evolutionary processes.

## Introduction

Tropical regions harbor most of the biodiversity that is encountered on earth and play a fundamental role in global environmental homeostasis [1,2]. Despite the fundamental role of tropical habitats, these remain severely understudied and particularly in the Neotropics, the biodiversity composition of many taxa still remains unknown [3]. High biodiversity ecosystems pose the challenge of species identification and delimitation, especially in areas where for many insects the last biodiversity survey was done at least a century ago, as is the case of fireflies [4,5]. Species delimitation methods [6,7] based on genetic information have proven to draw first evidence on formulating species delimitation hypothesis. An integrative species delimitation approach that additionally includes morphological, ecological and behavioral evidence would result in a confident analysis of the current status of a species, even though this approach can only be done for some species and at a slow processing rate.

The American continent has a complex geological and glaciation history that has shaped current biodiversity patterns [8,9]. Distribution patterns in the Americas have additional been influenced by the rise of the land bridge connecting South and North America ∼3-5 million years ago (mya) and more recently by the expansion and contraction of ice sheets during the last glaciation [10,11]. Nuclear Central America (NCA), from the Tehuantepec isthmus to the Nicaraguan depression, is a region harboring many different ecosystems types, including cloud forests, rain forests and subtropical thorn scrub habitats [9,12]. Additional to its classification as a biodiversity hotspot, NCA shows particularly high endemism levels where many taxa have speciated in the region [13,14]. As a consequence, NCA maintains a unique genetic diversity, making this region highly important for conservational efforts [15]. Fireflies are bioluminescent beetles with a world-wide distribution, with the exception of New Zealand and the polar caps [16]. The behavior and phylogenetic relationships of some firefly genera, especially from North America and Asia have been well studied, and have generated many insights into the evolution of bioluminescent behavior, their phylogenetic relationships and their biogeographical history [17–19]. On the other hand, very little is known about fireflies outside these regions, although research efforts are present [20–22].

In this work we collected fireflies in Guatemala and using a genetic barcoding approach, we generated time-calibrated phylogenetic trees focusing on the genera *Photinus, Photuris* and *Bicellonycha*. We used a Poisson Tree Process (mPTP) species delimitation model to generate species hypothesis of the collected fireflies. We also describe four new species of *Photinus* fireflies: *P. hunahpú, P. helodermensis, P. schusteri, P. semetabajense*. This works highlights the need for biodiversity research in the Neotropics and how narrowing sampling biases will improve our understanding on broad evolutionary processes in fireflies.

## Results

We classified 15 firefly morphospecies collected in different locations that include multiple ecosystem types (dry forest, cloud forest and pine/oak forest) (**Figure 1**). We generated time-calibrated phylogenetic trees separately for *Photinus* and *Photuris* together with *Bicellonycha* revealing the phylogenetic placement and divergence times of Guatemalan fireflies. The Guatemalan *Photinus* species diverged 28 mya (Oligocene) forming a distinct and unique clade from North American species (red node 1, **Figure 2**). After the Guatemalan and the North American clade diverged, *in situ* speciation and diversification took place at the end of the Miocene (∼ 18-21mya) and within species further diversification occurred in the Pliocene/Pleistocene.

**Figure 1.**
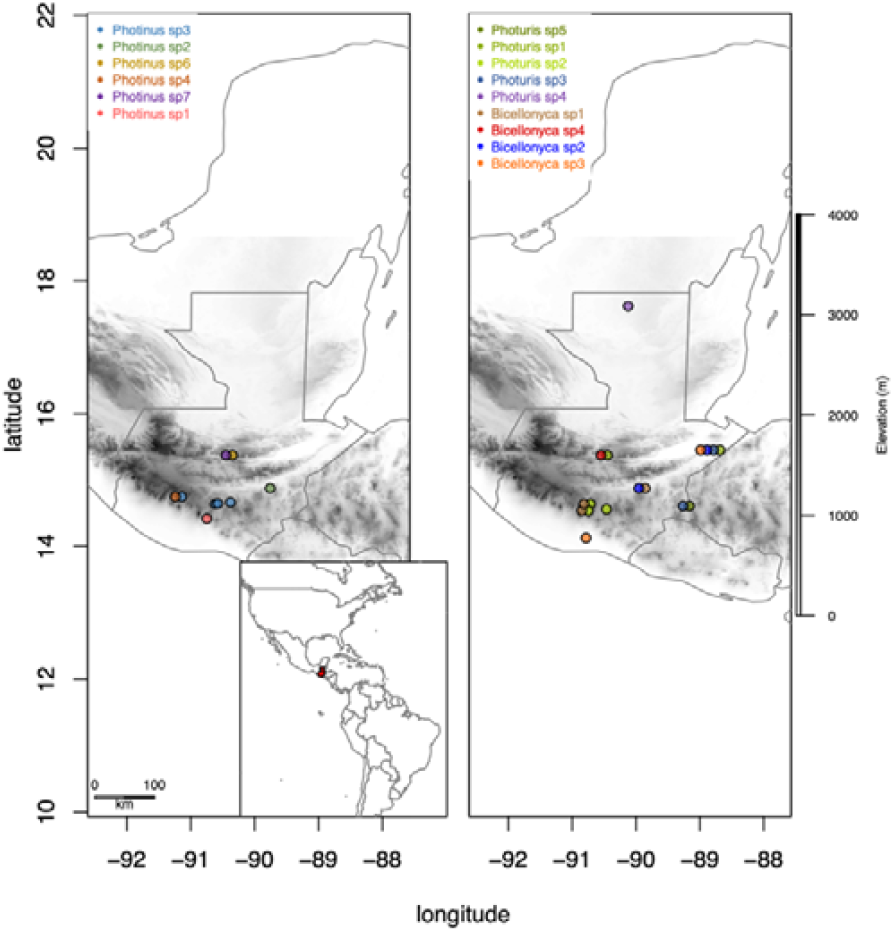
Map of Guatemala showing the distribution of collected *Photinus* (left panel), *Photuris* and *Bicellonycha* (right panel) fireflies. Colored dots in the map mark collecting sites for each of the species. Lateral bar indicates elevation level in meters. Inset: map of the American continent highlighting Guatemala in red.

**Figure 2.**
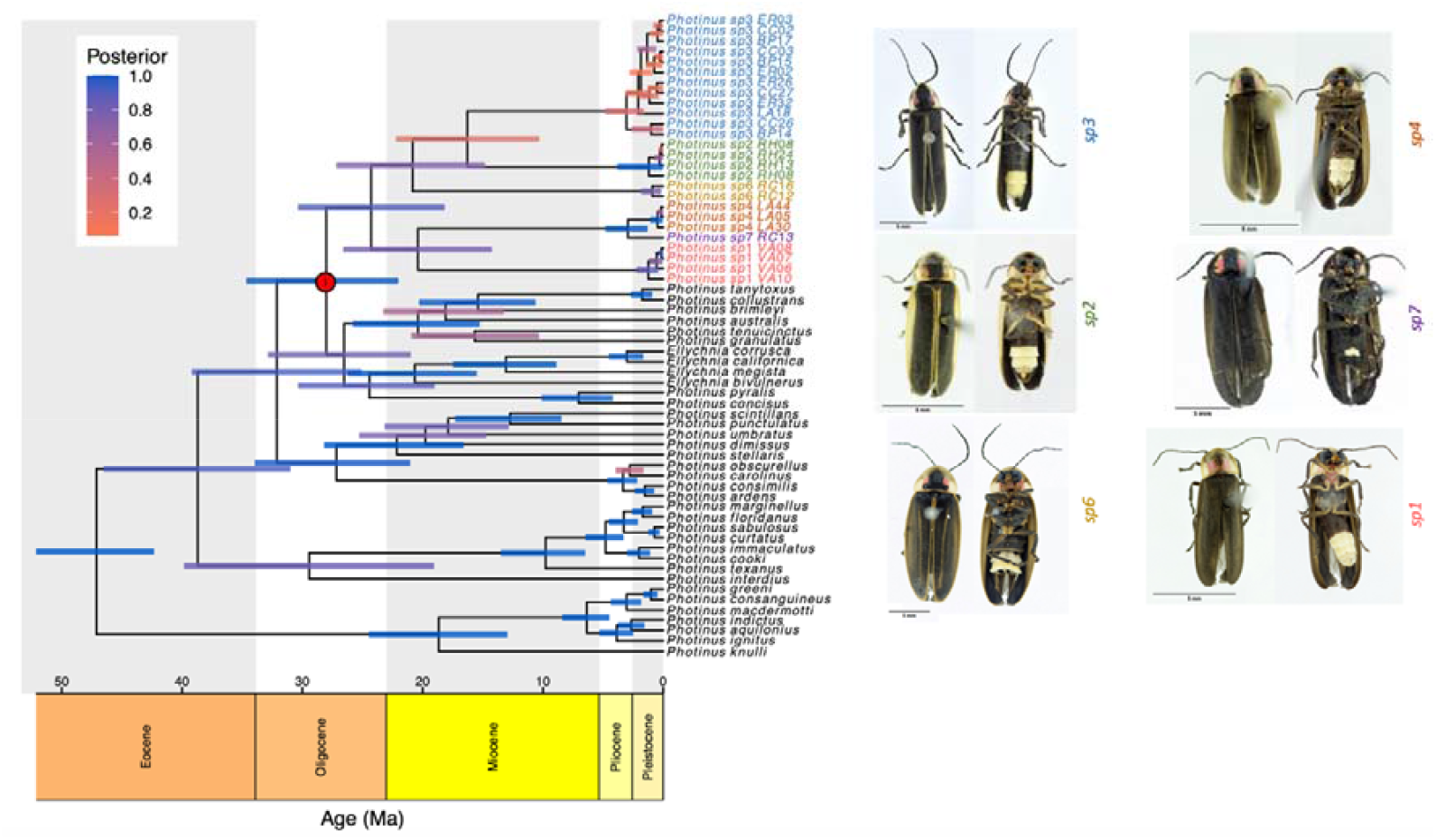
Time-calibrated phylogenetic tree of *Photinus* fireflies. Horizontal bars depict 95% credible intervals of the node’s ages. Label colors highlight defined species. Red dot highlights the divergent node between the Guatemalan and the North American *Photinus* node. Right panel: Images of the specimens placed in a phylogenetic context.

We used a multi-rate Poisson Tree Process (mPTP) to generate phylogeny-based species delimitation hypothesis using our time-calibrated tree [23]. We identified six Guatemalan *Photinus* species following the mPTP method which corresponded to the six identified morpho species (**Table S1**).

We performed morphological characterization of the collected *Photinus* by measuring several morphological traits (**Table S2**). A taxonomic assessment was done using existing taxonomic keys to fit the collected *Photinus* into already described species (see methods section). We also revised additional species descriptions and species lists for the region (see methods section) and together with the revision of *Photinus* type specimens, which included *P. picticollis, P. pulchellus*, and *P. affinis*, we propose four new species of *Photinus*: *Photinus hunahpú* (*Photinus sp1*), *Photinus helodermensis* (*Photinus sp2*), *Photinus schusteri* (*Photinus sp3*) and *Photinus semetabajense* (*Photinus sp4*). Taxonomic identification of *Photinus sp6* lead to placing this species as *Photinus congruus* (**Figure S5**) and for *Photinus sp7* (**Figure S6**) only two female individuals were collected. As there is no taxonomic information to key females, *Photinus sp7* was not able to be places into a species. A detailed description of the new *Photinus* species can be found in the supplementary materials (**Figures S1-S4**).

NCBI sequences of target loci were only present for eight *Photuris* and *Bicellonycha* species, as opposed to *Photinus*, where genetic information was found for 37 species. We classified five morphospecies for *Photuris* (**Figure S7-S11**) and four for *Bicellonycha* (**Figure S12-S15**). For *Bicellonycha* there is genetic information available only for three species [17] and only for *B. wickershamorum* sequence data for the loci of interest was found [16]. This reflects the lack of genetic studies for this genus and highlighting the need for biodiversity discovery research in fireflies. The addition of Nuclear Central American specimens reveals deeper divergent times between *Photuris* and *Bicellonycha* than previously observed, with the age for the oldest node for *Photuris* (**Figure 2**, node red - 1) placed at the early Oligocene ∼33 Mya. The *Photuris* node-1 diverges into two clades. Node 2, leads to a cluster including North American *Photuris* (*P. divisa* and *P. frontalis*) closely related to *Photuris sp4. Photuris sp4* (**Figure 1**, purple dot), was collected in the most northern part of Guatemala, at the Petén basin, which is part of the North American tectonic plate. Node 3, clusters only *Photuris* species reported to have a North American distribution [16]. Node 4, which dates at the beginning of the Pliocene ∼5.3Mya, leads to clusters with shorter branch lengths into the Pleistocene. Starting at the Pliocene, the Guatemalan *Photuris* clades show short branch lengths, hinting at fast diversification in the region.

The oldest node for *Bicellonycha* was placed at the end of the Eocene, ∼ 36 Mya (**Figure 3**, node - 5). Our sampled *Bicellonycha* formed two clades that further diverged into the Miocene and which reflects deep phylogenetic relationships within *Bicellonycha*. The similar external morphology between *Bicellonycha sp2* and *Bicellonycha sp3* but deep divergence times, poses a challenge for species identification using external morphological traits and increases the chance of the presence of cryptic species within this genus. Species delimitation inference using mPTP proposed three *Photuris* species out of the five that we identified as morphospecies (**Figure 3, Table S1**), clustering morphospecies 1, 2 and 5 into a single species. Taxonomic key species identification could not place the collected Guatemalan *Photuris* into already described species [22]. Our morphospecies classification of *Bicellonyhca* is in concordance with the mPTP proposed species, identifying four species.

**Figure 3.**
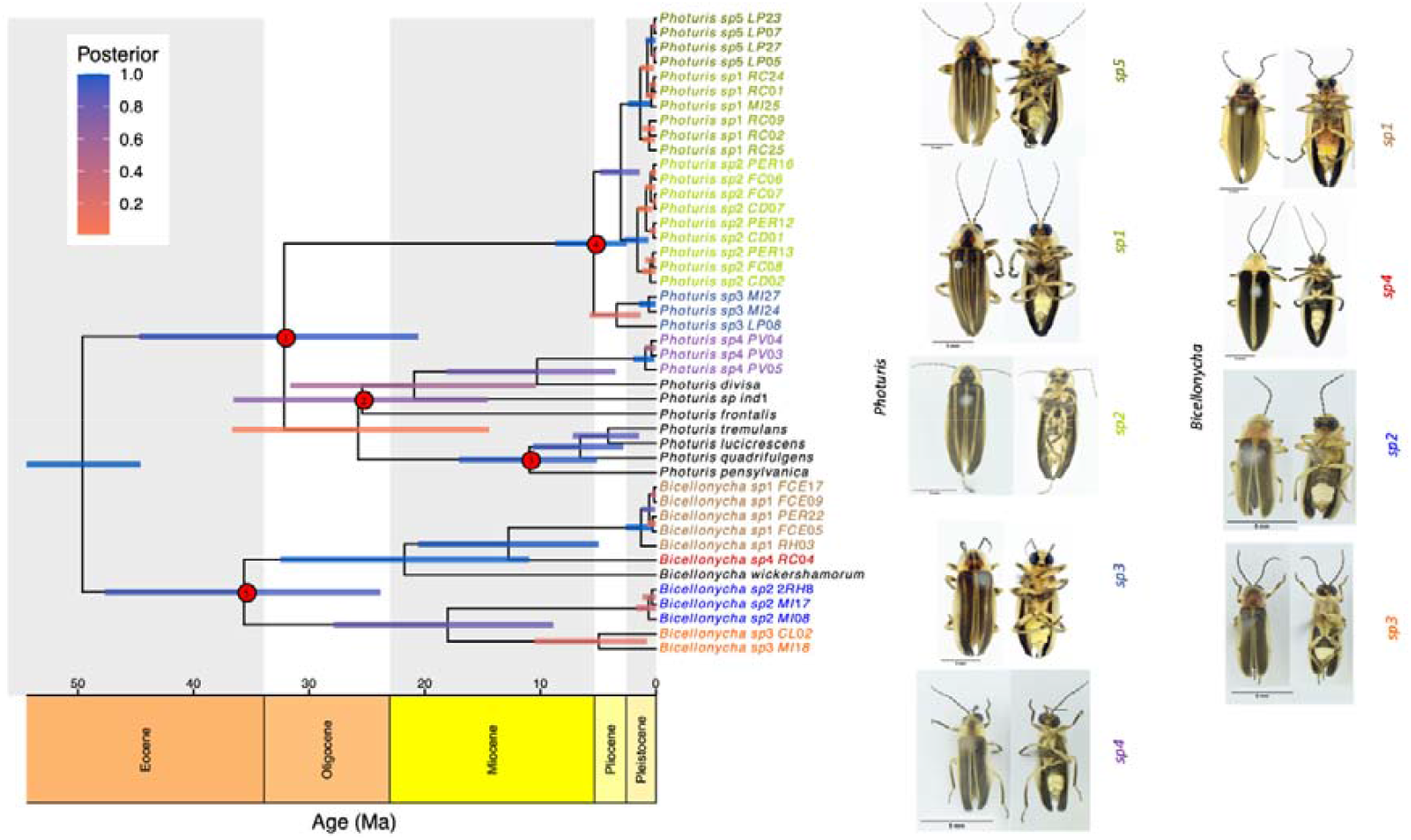
Calibrated phylogenetic tree of *Photuris* and *Bicellonycha* fireflies. Horizontal bars depict posterior probability of the node’s ages. Label colors highlight putative morphospecies. Red dots highlight the nodes leading to species collected in Guatemala. Right panel: Images of the specimens placed in a phylogenetic context.

## Discussion

Research on biodiversity resources is fundamental for the conservation of biomes and the most basic stepping-stone for the understanding of the ecology and evolutionary history of species. Biodiversity research in Central American Neotropical fireflies is scarce. The last biodiversity/taxonomic survey was done with the Biologia Centrali Americana in 1881 [24] and since then, limited efforts have been published on the study of firefly biodiversity in Nuclear Central America [25,26]. This work presents the first overview on the phylogenetic placement of Guatemalan fireflies and their divergence times. We also describe four new species for *Photinus*: *P. hunahpú, P. helodermensis, P. schusteri, P. semetabajense*. Our *Photinus* were collected in different types of ecosystems, including; low land coastal tropical forest (*P. hunahpú*), dry shrub forest (*P. helodermensis*), highland pine-tree forests (*P. schusteri* and *P. semetabajense*) and cloud forest (*Photinus sp5* and *Photinus sp7*). These distinctive ecosystems together with our species delimitation analyses suggest that our four new *Photinus* firefly species are endemic to Guatemala. The high diversity found in Nuclear Central America underlines the high potential for species discovery in this region.

*Photinus* is a species diverse genus, with a broad distribution in the Americas (Canada to Argentina) occupying a wide range of ecosystems [27]. Our morphospecies assignment for the presented *Photinus* was in concordance with the species delimitation results obtained by mPTP, identifying a total of seven species (**Figure 2**). Future genetic and morphological information gathered from its range distribution will then shed light into the evolutionary processes defining *Photinus* diversity and distribution.

*Photuris* species have been reported to feed on other fireflies, a behavior that is reflected in their external morphology [28,29]. Research on the behavior of Central American *Photuris* is pending, but external morphological traits such as strong mandibles, thorax and legs, suggest that our *Photuris* species might also show a predatory behavior. Morphospecies *Photuris sp1, Photuris sp2* and *Photuris sp5* show very similar but not identical morphology. From a genetic perspective, these three species were pooled as one putative species by the mPTP model (**Figure 3**). *Photuris sp1-2-5*, species complex has a broad distribution (**Figure 1**) as has been reported for other *Photuris* species [28]. Species with a wide distribution range tend to show phenotypic variation due to natural selection or drift [30,31]. Therefore, morphologic variation across populations cannot be ruled. Overall, there is less genetic, behavioral and ecological information about *Bicellonycha* [26,32–34]. Here we present the phylogenetic relationships and divergent times of four *Bicellonycha* species revealing a clade with deep divergent times (**Figure 3**). This genus seems to be widespread distributed in Guatemala, occurring in tropical, cloud and dry forest as well as in coastal ecosystems (**Figure 1**).

*Photinus, Photuris* and *Bicellonycha* are distributed in the whole American continent [35,36]. Most of the gathered genetic data for these genera comes mostly from sampled data in North America [17,18]. The deep divergent times observed between the Guatemalan and the North American species in the *Photinus* and *Photuris*/*Bicellonycha* trees, opens up questions about its biogeographic history. Our phylogenetic results suggest a single colonialization of *Photinus* in Guatemala as all Guatemala specimen formed a monophyletic clade (Figure 2). The biogeographic history of *Photuris* and *Bicellonycha* appears to be more complex with potentially multiple colonialization events, as the Guatemalan specimens did not form a monophyletic clade with respect to the North American of *Photuris* and *Bicellonycha* (Figure 3). However, it could be possible that *Photuris* and *Bicellonycha* colonialized North America in a single event. Collecting more specimens from different geographical regions in the American continent will help to uncover past colonization processes as well as speciation and ration events. Work on Nuclear Central American fireflies shows a high potential for species discovery, as well as new reports for endemic species, as this region is characteristic for its high endemism [37,38].

## Conclusions

Species delimitation is a challenging task and leveraging genetic and morphological information can produce well founded hypotheses on the status of species. Factors such as gene flow between populations can generate insights into populations connectivity, information that might be crucial to understanding the process of speciation and the morphological diversity observed. Differences in nucleotide diversity levels within species could also be informative for the definition of genetic groups and approaches such as polymorphism aware models (PoMos) have the potential to use polymorphic information to define species [39]. In the case of flashing fireflies, information on the flashing pattern (flash frequency, duration, flying height) has been reported to be useful for the identification of species [25,40] and this axis of evidence would further provide useful information for species delimitation.

## Methods

### Sample collection and lab work

Sample collection was done at 13 locations (Figure1 and **Table S3**) using an insect net. Specimens were stored in 96% ethanol and kept at -20°C. DNA extractions were done using the Promega Wizard Genomic DNA Purification kit, following the manufacturer’s guidelines and DNA quality and concentration was measured using a Nanodrop.

One mitochondrial gene (COI) and two nuclear genes (28s and 18s) were amplified using polymerase chain reaction (PCR) with primers and amplification profiles described in **Table S4**. PCR reactions having the expected gel electrophoresis band size were send for Sanger sequencing at Macrogene.

### Species diagnosis and description

We conducted a morphometric study for the taxonomic identification of the collected specimens. For this we revised reported species for Guatemala described in the Biologia Centrali Americana (BCA) [24], The Coleopterorum Catalogus [41], Checklist of the Coleopterus Insects of Mexico, Central America, the West Indies and South America [42], and the document, Materials for a revision of the Lampyridae [43]. We also keyed the specimens using the following taxonomic keys: La familia Lampyridae (Coleoptera) en la estación de biología tropical “Los Tuxtlas”, Veracruz, Mexico [44], The Fireflies of Ontario [45], Luciérnagas del centro de México (Coleoptera: Lampyridae): descripción de 37 especies nuevas [22] and Luciérnagas de la región golfo-Caribe de México y descripción de 16 especies nuevas [46]. Specifically for *Photinus*, type specimens from the region were imaged by the Natural History Museum of London, providing dorsal and ventral images of *Photinus picticollis, Photinus pulchellus* and *Photinus affinis* [<u>link 1</u>].

Measurements in millimeters were taken with a Wild M3B Typ 308700 stereoscope. A standardized graticule was used to take the following morphometric measurements (between parentheses is explained how they were measured): total length (from the apex of the pronotum to the base of the elytra), elytral width (on the right side, over the humeral space), length of elytra (on the right side), pronotum length (over the middle part of the pronotum), pronotum width (measured at the widest part of the pronotum), antenna length (it was calculated as the sum of the length of each separate antennomere; starting from the proximal antennomere), head width (dorsal side, was taken from where an eye begins to where the eye ends), head length (dorsal side, was taken from where the head is visible on the dorsal side to where the jaws start), eye width (right side, measured in side view), eye length (right side, measured in side view), length of maxillary palps (calculated as the sum of the length of the four separate maxillary palp parts; starting from the proximal palp), interantenal fossae width (right side, measured in the middle part of the interantenal fossae), interantenal distance (measured in the middle part of the interantenal space), interocular distance (dorsal side), length of sternites from 1-7 (measurement was taken at the right end of each sternum).

Aedeagus were dissected for each individual, mounted on cardboard sheets and placed on the mounting pins of the corresponding specimens. Photographs were taken with a Nikon D7200 and a 105mm lens. The species have been described following the general terminology [22].

### Bioinformatic analysis

Sanger sequencing traces were curated using MegaX [47]. Published sequences were retrieved from NCBI using Biopython by implementing the tags “Lampyridae”, “COI”, “18s” and “28s”. All fasta files were aligned using AliView [48] and were manually curated. For our phylogenetic analysis of *Photinus* we additionally used the previous alignments of the CAD, 16s, UV opsin and WG genes to improve phylogenetic resolution [49]. A phylogenetic analysis was performed using the software RevBayes [50]. Specifically, we applied a by-locus partitioned phylogenetic model with independent rate-multipliers per locus and independent substitution model parameters. That is, we assumed an independent GTR+Gamma substitution for each locus. Since insufficient fossil information is available to perform a primary calibration, we used the estimated ages of *Photinus* and *Photinus+Bicellonycha* from Catalán et al 2024 [51] as secondary calibrations. Specifically, we applied a root node calibration using a normal distribution with mean 47.5 and standard deviation 2.5 for Photinus and a normal distribution with mean 50 and standard deviation of 2.5 for *Photinus+Bicellonycha*. We applied an uncorrelated lognormal relaxed clock model with estimated mean and standard deviation. We ran 8 replicated Markov chain Monte Carlo analyses with 500,000 iterations, sampling every 10^th^ iteration. We checked for convergence using the R package Convenience [52]. The posterior sample of phylogenies was summarized as the maximum a posteriori phylogeny, i.e., the topology with the highest posterior probability. To generate species delimitation hypothesis we used the mPTP method [7]. The following command was used to generate species delimitation hypothesis: ‘mptp --mcmc 50000000 --multi --mcmc_sample 1000000 --mcmc_burnin 1000000’. Generated fasta files for each loci and calibrated trees were uploaded to dryad DOI: 10.5061/dryad.66t1g1kbr

## Author’s contributions

AC conceived the study and wrote the manuscript with input of CMPA, AMEM, JAGG, VVB and SH. CMPA, AMEM, JAGG and did sample collection, molecular work and sequence curation. VVB made the species descriptions and phylogenetic based analyses were done by AC and SH.

## Acknowledgments

We would like to thank Gabriela Alfaro and the biology department at UVG for administrative support, to Alejandra Zamora for her advice and support in the molecular lab and to xx for support in sample collection. To Michael Geiser and Keita Matsumoto from Natural History Museum of London for providing images of type specimens. This work was founded DFG SPP-1991 to SH and AC.

