## Supplementary material for "Filling the gap in the Neotropics: new firefly species provide insights into Central American biodiversity": Suplementary file

### Taxonomic descriptions of four *Photinus* species.

Lampyridae: Lampyrinae: Photinini

***Photinus hunahpu*** **(*Photinus sp1*) Catalán et al, sp. nov.**
**Figure S1**

**Diagnosis.** Body 7.50-9.00 long and 2.40-2.70 wide, antennae filiform, twice the length of pronotum, robust mandibles and terminal maxillary palpomere securiform; pronotum wider than long, with two reddish spots; elytra parallel, more than five times longer than wide; claws simple; aedeagus robust, with broad basal piece, two dark-colored excrescences protruding from the middle lobe, dorsal part slightly sclerotized and ventral part membranous.

**Description**. Male. Body length 9.00; maximum body width 2.40 (pronotum). Body small, ovoid, brown, except for the pronotum (with two reddish spots), scutellum, lateral and sutural margin of elytra, pro- and mesothorax dark brown to black; trochanters and basal part of femora yellow. The luminous apparatus occupies the entire surface of sternites 5-6 and one third of sternite 4. **Head**. Interocular distance (0.50) concave, shiny and sparsely pilose; forehead vertical; interantennal distance (0.07) half the width of antennal fossa (0.14 ); eyes finely faceted, spherical, as long (0.96) as wide (0.96), almost twice the size of the interocular distance; antennae filiform, 2-2.5 times the length of pronotum, surpassing mesocoxae, with dense setae, antennomere 1 claviform (0.50) shorter than the next two combined; frontoclipeal suture membranous, concave; clipeus trapezoid-shaped, posterior margin curved with middle notch, surface with long, regularly distributed setae; mandibles robust, with few external setae and ending in a sharp stylet; labellum membranous; terminal maxillary palpomere securiform (0.29), twice as long as the preceding three combined (0.76); terminal labial palpomere securiform. **Thorax.** Pronotum wider (2.58) than long (1.80), semicircular, posterior margin straight, posterior angles acute, disc concave, sides broadly flattened, surface shiny, porous, with a long and abundant decumbent pilosity; scutellum triangular, surface flat and pilose, with posterior margin rounded; elytra parallel, more than five times longer (7.05) than wide (1.20), surface roughly dotted, shiny and with decumbent pilosity; mesothoracic respiratory spiracles tubular; legs similar to each other, femur fusiform, flattened, tibiae flattened, outer margin entire, tarsomeres laterally not compressed, tarsomere 1 (0.36) half the length of tarsomere 2, one third of tarsomere 3, similar to tarsomeres 4 and 5, claws simple. **Abdomen.** With 8 visible sternites, sternites 1-4 of similar length (0.54-0. 60), sternites 5-6 similar to above, with distinct stigmatiform pores, sternites 5-6 with margin notched, sternite 7 with margin concave; sternite 8 with margin bullet-shaped; posterior margin of pygidium rounded; aedeagus robust, S-shaped, with broad basal piece, two dark-colored excrescences protruding from the middle lobe, parameres rounded, almost completely covering the middle lobe dorsally, with obtuse apex, dorsal part slightly sclerotized and ventral part membranous. Variation: 7.50 to 9.00 long and 2.40 to 2.7 width.

**Etymology.** Specimens of this species were collected at the Agua volcano (Volcán de Agua), previously known by the Kaqchikel Mayan communities as Hunahpú. Hunahpú is also the name of one of the Mayan hero twins depicted in the Popol Vuh.

**Material examined:**

***Holotype***, male. GUATEMALA: Palín, Escuintla, south side of the Agua volcano [14.414657,-90.748566]. Collecting date: 04-09-2021. Col.: Adriana Echeverría.

***Paratypes*** (n=3), two males, one female. GUATEMALA: Palín, Escuintla, south side of the Agua volcano [14.414657,-90.748566]. Collecting date: 04-09-2021. Col.: Adriana Echeverría.

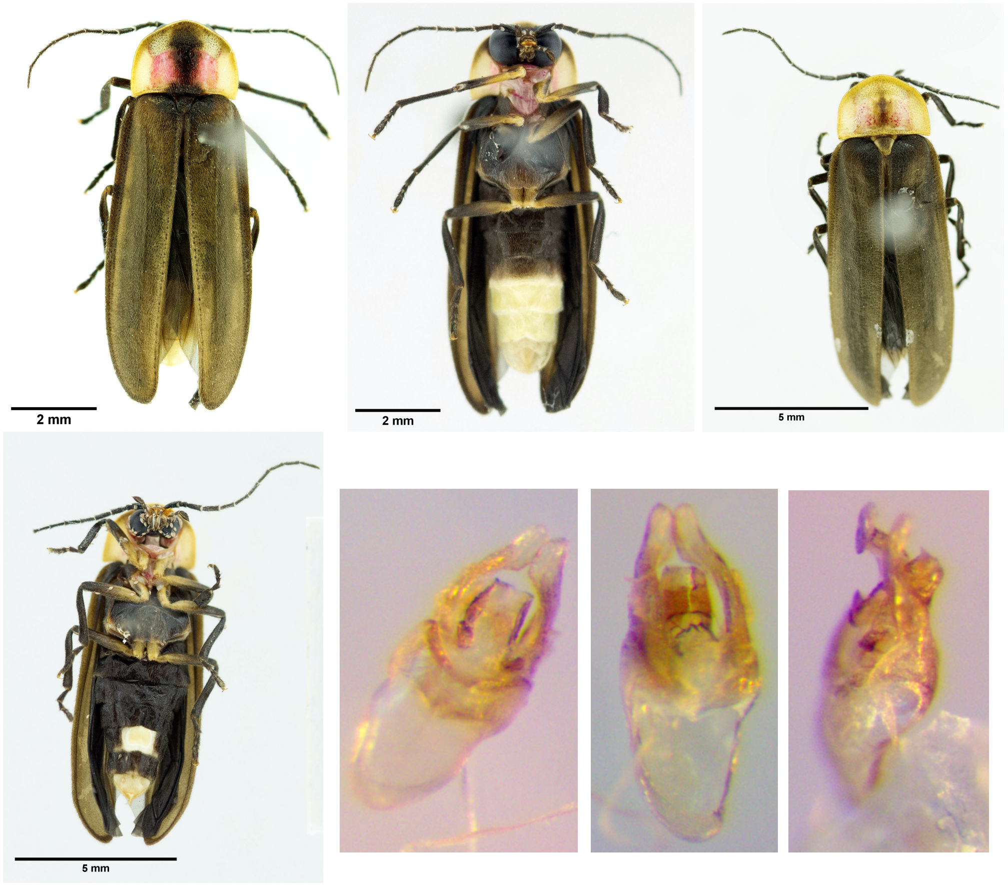


**Figure S1.** *Photinus hunahpu*. Upper panel: male dorsal and ventral view. Female dorsal view. Lower panel: female ventral virew. *Edeagos*: dorsal, ventral and lateral views.

***Photinus helodermensis* (*Photinus sp2*) Catalán et al, sp. nov.**
**Figures S2**
**Diagnosis.** Body 8.10-10.05 long and 2.10-3.00 wide, antennae filiform, twice the length of pronotum; semicircular pronotum, wider than long; elytra ovoid, 6 times longer than wide, surface roughly dotted, shiny and with decumbent pilosity; claws simple; aedeagus robust, wide basal piece, two dark-colored excrescences protruding from the middle lobe, dorsal part slightly sclerotized and ventral part membranous.
**Description**. Male. Body length 8.10; maximum body width 2.10 (pronotum). Body small, oblong, pro- and mesothorax dark brown to black except or the pronotum (with two reddish spots), lateral and sutural margin of elytra pale yellow, trochanters and basal part of femora yellowish. The luminous apparatus occupies the entire surface of sternites 5-6. **Head**. Interocular distance (0.60 ) excavated, shiny and sparsely pilose; forehead vertical; interantennal distance (0.10) half the width of antennal fossa (0.21); eyes finely faceted, spherical, as long (0.90) as wide (0.90), less than twice as long as the interocular distance; antennae filiform, twice the length of pronotum, surpassing mesocoxae, with dense setae, antennomere 1 claviform (0.57) about the same length as the next two combined; frontoclipeal suture membranous, concave; clipeus trapezoid-shaped, posterior margin straight, surface with long, regularly distributed setae; mandibles robust, with few external setae and ending in a sharp stylet; labellum membranous; terminal maxillary palpomere securiform (0.31), 2.5 times longer than the preceding three combined (0.8); terminal labial palpomere securiform. **Thorax.** Pronotum wider (2.40) than long (1.74 ), semicircular, posterior margin straight, posterior angles acute, disc concave, sides broadly flattened, surface shiny, porous, with a long and abundant decumbent pilosity; scutellum triangular, surface

concave and pilose, with posterior margin rounded; elytra ovoid, 6 times longer (6.45 ) than wide (1.05), surface roughly dotted, shiny and with decumbent pilosity; mesothoracic respiratory spiracles tubular; legs similar to each other, femur fusiform, flattened, tibiae flattened,with a wider apex, outer margin entire, tarsomeres laterally compressed, tarsomere 1 (0.30) similar in length to tarsomeres 2 and 4, one third of tarsomeres 3 and 5, single claws. **Abdomen:** With 8 visible sternites, sternites 1-2 of similar length (0.54 mm), sternite 4 shorter (0.42), sternites 3, 5 and 6 similar (0.60), with distinct stigmatiform pores, sternites 5-6 with margin with a small median notch, sternite 7 with margin with a strongly notch; sternite 8 with margin bullet-shaped; posterior margin of pygidium rounded; aedeagus robust, S-shaped, with a wide basal piece, two dark-colored excrescences protruding from the middle lobe, parameres rounded, almost completely covering the middle lobe dorsally, with obtuse apex, dorsal part slightly sclerotized and ventral part membranous. Variation: 8.10 mm to 10.05 mm long and 2.10 to 3.00 width.

**Etymology.** This is the first firefly species described from the Guatemalan dry shrub forest located at the department of Zacapa. The holotype and paratype specimens were collected at the Heloderma Natural Reserve, which was founded to conserve this unique ecosystem.

**Material examined:**

***Holotype***, male. GUATEMALA: Reserva Natural del Heloderma, El Arenal, Zacapa. [14.8738028,-89.7569861]. Collecting date: 23-07-2021. Col.: José Andrés Gutiérrez.

***Paratypes*** (n=6). Males. GUATEMALA: Reserva Natural del Heloderma, El Arenal, Zacapa. [14.8738028,-89.7569861]. Collecting date: 23/24-07-2021. Col.: José Andrés Gutiérrez.

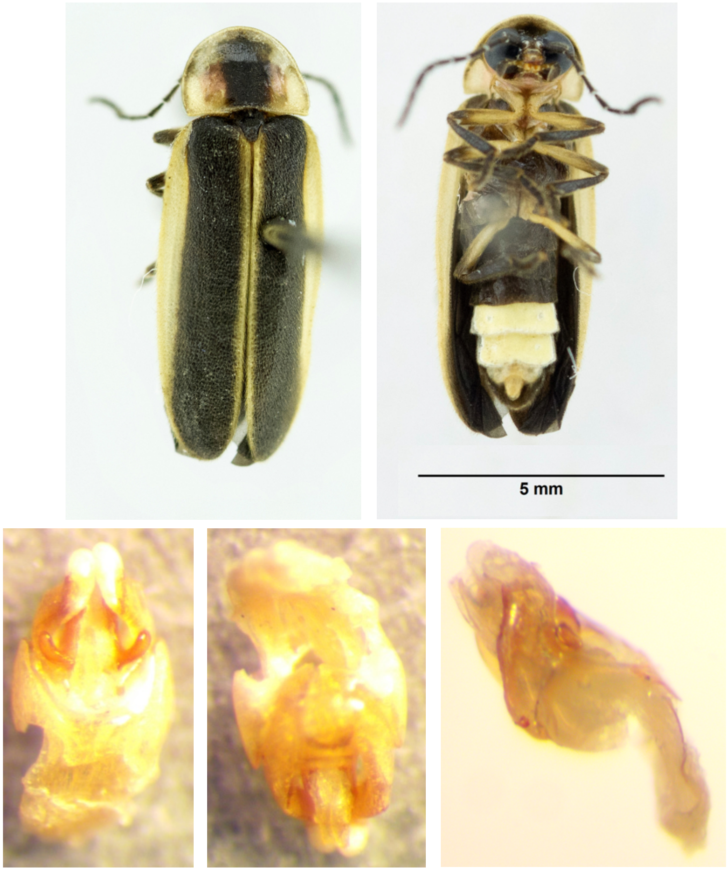


**Figure S2.** *Photinus helodermensis*. Upper panel: male dorsal and ventral view. Lower panel: *Edeagos*: dorsal, ventral and lateral views.

***Photinus schusteri*** **(*Photinus sp3*) Catalán et al, sp. nov.**
**Figures S3**
**Diagnosis.** Body 10.80-12.90 long and 2.40-3.30 wide, parallel, black; the luminous apparatus occupies the entire surface of sternites 5-6 and one third of sternite 4; antennae filiform, slightly serrate, twice the length of pronotum; bullet-shaped pronotum with two reddish spots, wider than long; elytra ovoid, 6 times longer than wide, surface roughly dotted, shiny and with decumbent pilosity; claws simple; aedeagus robust, wide basal piece, two dark-colored excrescences protruding from the middle lobe, dorsal part slightly sclerotized and ventral part membranous.
**Description**. Male. Body length 12.90; maximum body width 3.30 (pronotum). Body long, parallel, black, except for the pronotum (with two reddish spots), lateral and sutural margin of elytra pale yellow, pro- and mesothorax dark brown to black, trochanters and basal part of femora yellow. The luminous apparatus occupies the entire surface of sternites 5-6 and one third of sternite 4. **Head**. Interocular distance (0.86) concave, shiny and sparsely pilose; forehead vertical; interantennal distance (0.24) as long as antennal fossa (0.29 mm); eyes finely faceted, spherical, as long (1.14 ) as wide (1.14 ), but less than twice as long as the interocular distance; antennae filiform, slightly serrate, twice the length of pronotum, surpassing mesocoxae, with dense setae, antennomere 1claviform (0.67) shorter than the next two combined; frontoclipeal suture membranous, concave; clipeus trapezoid-shaped, posterior margin curved, surface with long, regularly distributed setae; mandibles robust, with few external setae; labellum membranous; terminal maxillary palpomere securiform (0.38), twice as long as the preceding three combined and three times more than the proximal; terminal labial palpomere spindle-shaped. **Thorax.** Pronotum wider (3.12 mm) than long (2.64 mm), bullet-shaped, posterior margin straight, posterior angles acute, disc convex, sides broadly flattened, surface shiny, porous, with a small decumbent pilosity; scutellum triangular, surface concave and pilose, with posterior margin rounded; elytra parallel, more than five times longer (10.35 mm) than wide (1.65), surface roughly dotted, shiny and with decumbent pilosity; mesothoracic respiratory spiracles membranous; legs similar to each other, femur fusiform, flattened, tibiae flattened, with a wider apex, outer margin entire, tarsomeres laterally compressed, tarsomere 1 (0.54) almost twice as long as tarsomeres 2, 3, 4 and 5, claws simple. **Abdomen.** With 8 visible sternites, sternites 2-4 of similar length (0.84 ), sternites 5-6 longer than the previous ones (1.08 mm), and as long as sternite 1, with distinct stigmatiform pores, sternites 5-6 with margin with a small median notch, sternite 7 with margin concave; sternite 8 with margin bullet-shaped; posterior margin of pygidium acute; aedeagus robust, S-shaped, with a wide basal piece, and the posterior border showing a obtuse angled shape, two dark-colored excrescences protruding from the middle lobe, parameres rounded, almost completely covering the middle lobe dorsally, with obtuse apex, dorsal part sclerotized and ventral part membranous, terminal orifice. Variation: 10.80 to 12.90 long and 2.40 to 3.30 width.
**Etymology.** This species is dedicated to the entomologist Jack C. Schuster who significantly contributed to our knowledge on Guatemalan beetle biodiversity, including lampyrids.
**Material examined:**
***Holotype***, male. GUATEMALA: Zona 3, Mixco, Nueva Montserrat, Guatemala City. [14.642556, -90.573236]. Collecting date: 23-05-2021. Col.: Claudia Pérez.
***Paratypes*** (n=3). Males. GUATEMALA: Zona 3, Mixco, Nueva Montserrat, Guatemala City. [14.642556, -90.573236]. Collecting date: 23-05-2021. Col.: Claudia Pérez.

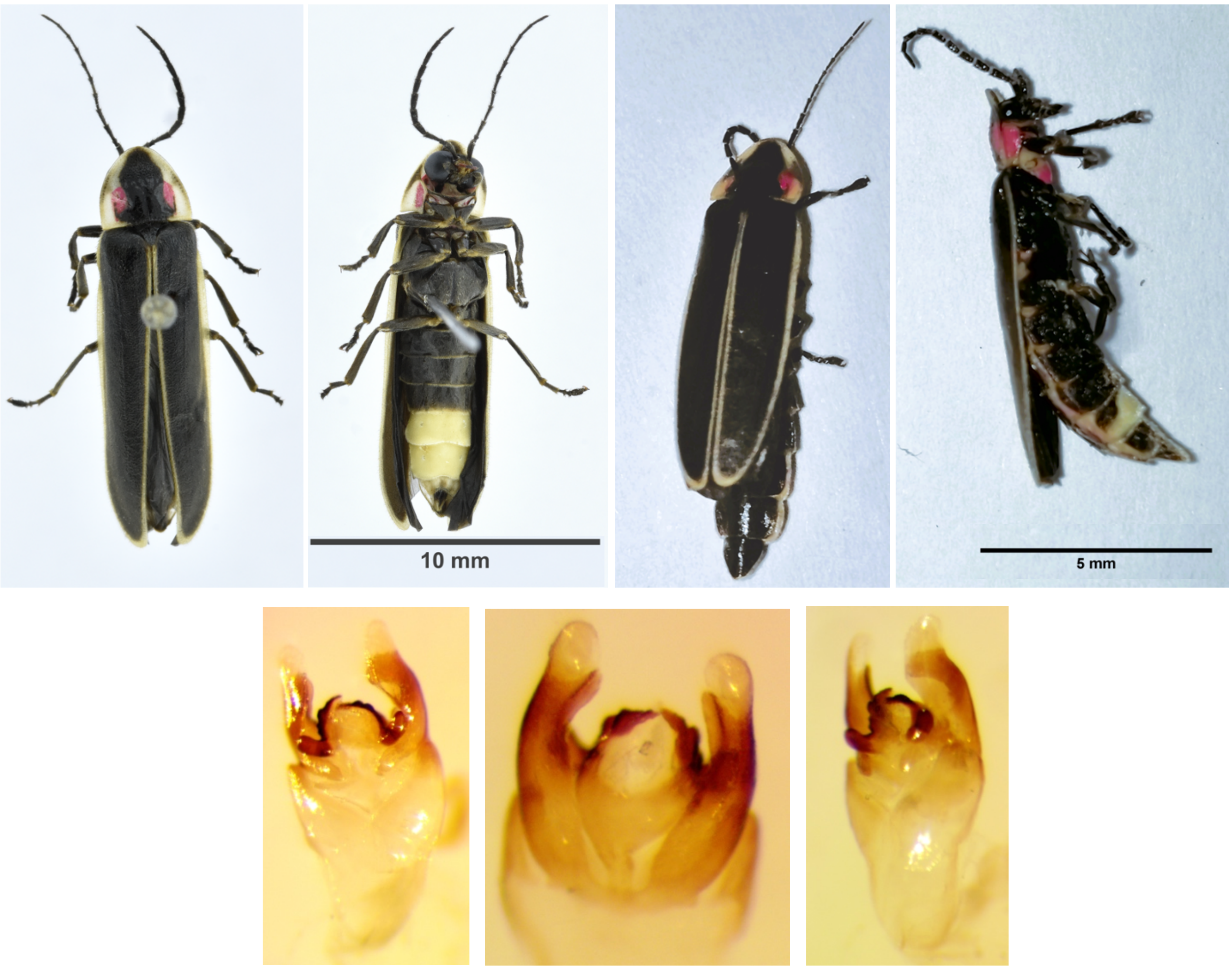


**Figure S3.** *Photinus schusteri.* Upper panel: *male* dorsal and ventral view. Female dorsal and ventral view. Lower panel: Edeagos: dorsal, ventral and lateral views.

***Photinus semetabajense*** **(*Photinus sp4*) Catalán et al, sp. nov.**
**Figures S4**
**Diagnosis.** Small body, 6.45-7.65 long and 2.10-.40 wide, ovoid darker brown; antennae filiform, twice the length of pronotum; semicircular pronotum with posterior margin sinuate, pale yellow with two reddish spots, wider than long; elytra semi-parallel, 5.5-5.7 times longer than wide; claws simple; sternites 5-6 longer than the previous ones with distinct stigmatiform pores and margin notched; aedeagus robust, wide basal piece, two dark-colored excrescences protruding from the middle lobe, dorsal part slightly sclerotized and ventral part slightly sclerotized.
**Description**. Male. Body length 7.35; maximum body width 2.10 (pronotum). Body small, ovoid darker brown, except for the pronotum pale yellow (with two reddish spots), pro- and mesothorax, trochanters and basal part of femora dark brown. The luminous apparatus occupies the entire surface of sternites 5-6 and one third of sternite 7. **Head**. Interocular distance (0.40) concave, shiny and sparsely pilose; forehead vertical; interantennal distance (0.07) half the width of antennal fossa (0.12 mm); eyes finely faceted, spherical, as long (0.72) as wide (0.72), almost twice as long as the interocular distance; antennae filiform, twice the length of pronotum, surpassing mesocoxae, with dense setae, antennomere 1 claviform (0.45 mm) longer than the next two combined; frontoclipeal suture membranous, concave; clipeus trapezoid-shaped, posterior margin curved, surface with long, regularly distributed setae; mandibles thin, with few external setae and ending in a sharp stylet;

labellum membranous; terminal maxillary palpomere securiform (0.21 ), twice as long as the preceding three combined; terminal labial palpomere securiform. **Thorax.** Pronotum wider (2.04 ) than long (1.44 ), semicircular, posterior margin sinuate, posterior angles acute, disc flat, sides broadly flattened, surface shiny, porous, with a small and decumbent pilosity; scutellum triangular, surface concave and pilose, with posterior margin rounded; elytra semi-parallel, 5.5 and 5.7 times longer (6.00) than wide (1.05), surface roughly dotted, opaque and with decumbent pilosity; mesothoracic respiratory spiracles membranous; legs similar to each other, femur fusiform, flattened, tibiae flattened, with a wider apex, outer margin entire, tarsomeres laterally not compressed, tarsomere 1 (0.24) half the length of tarsomere 2, tarsomeres 4 and 5 almost four times longer than tarsomere 3, claws simple. **Abdomen.** With 8 visible sternites, sternites 1 and 4 of similar length (0.48 ), sternites 5-6 longer than the previous ones (0.54-0.60), and as long as sternites 2-3, with distinct stigmatiform pores, sternites 5-6 with margin notched, sternite 7 with margin concave; sternite 8 with margin bullet-shaped; posterior margin of pygidium acute; dorsally open abdominal spiracles; aedeagus robust, S-shaped, with broad basal piece, the posterior border showing an obtuse angle, two dark-colored excrescences protruding from the middle lobe, parameres rounded, completely covering the middle lobe dorsally, with obtuse apex, dorsal part sclerotized and ventral part slightly sclerotized. Variation: 6.45 to 7.65 long and 2.10 to 2.40 width.
**Etymology.** The holotype and paratypes specimens were collected in the town of San Andrés Semetabaj, Sololá, therefore the specific epithet for *P. semetabajense* alludes to its collecting site.

**Material examined:**

***Holotype***, male. GUATEMALA: Lomas de Atitlán, San Andrés Semetabaj, Sololá [14.7444, -91.1386]. Collecting date: 26-06-2021. Col.: José Andrés Gutiérrez.

***Paratypes*** (n=3). Males. GUATEMALA: Lomas de Atitlán, San Andrés Semetabaj, Sololá [14.7444, -91.1386]. Collecting date: 26-06-2021. Col.: José Andrés Gutiérrez.

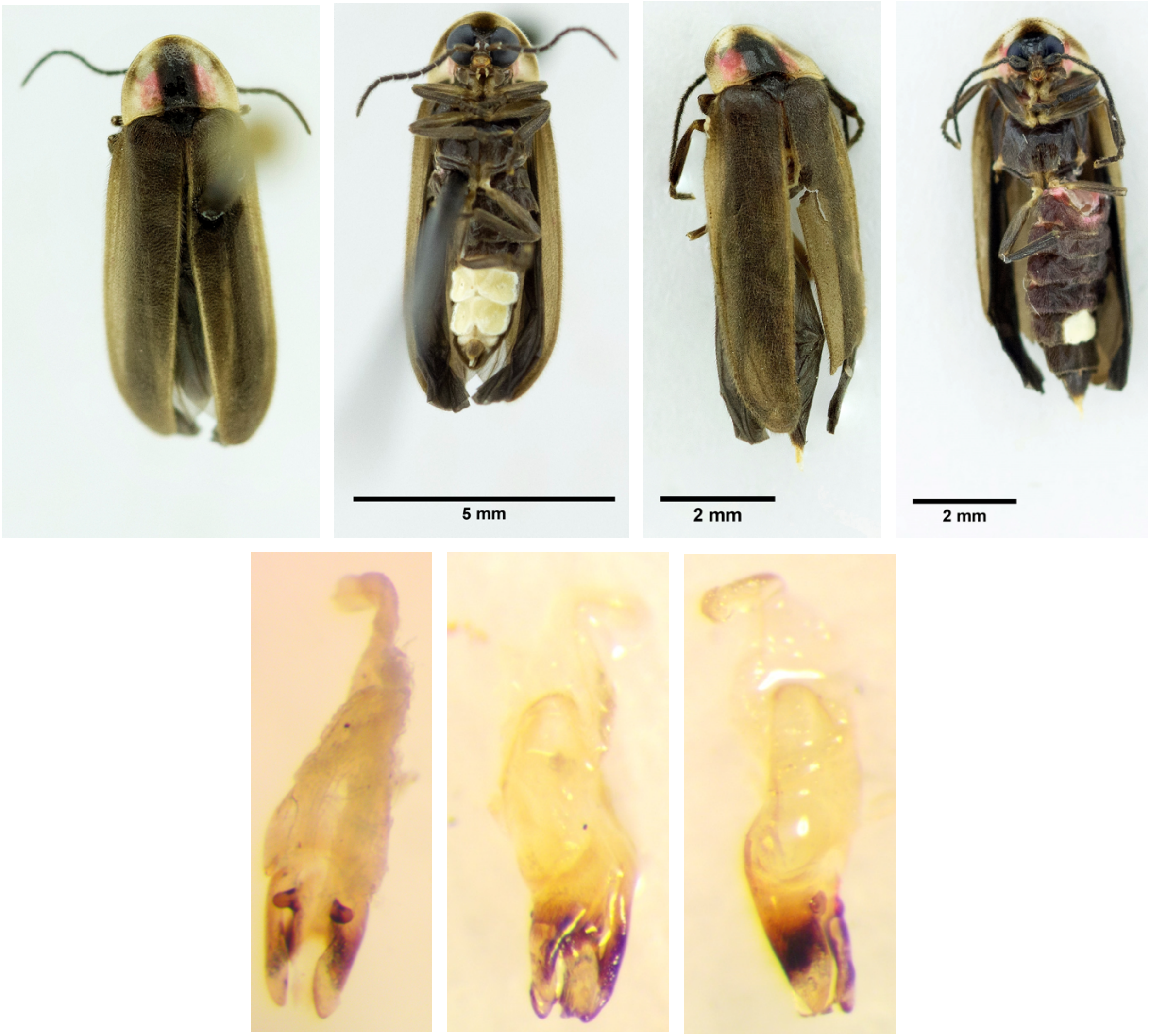


**Figure S4.** *Photinus semetabajense.* Upper panel: *male* dorsal and ventral view. Female dorsal and ventral view. Lower panel: Edeagos: dorsal, ventral and lateral views.

***Photinus congruus***

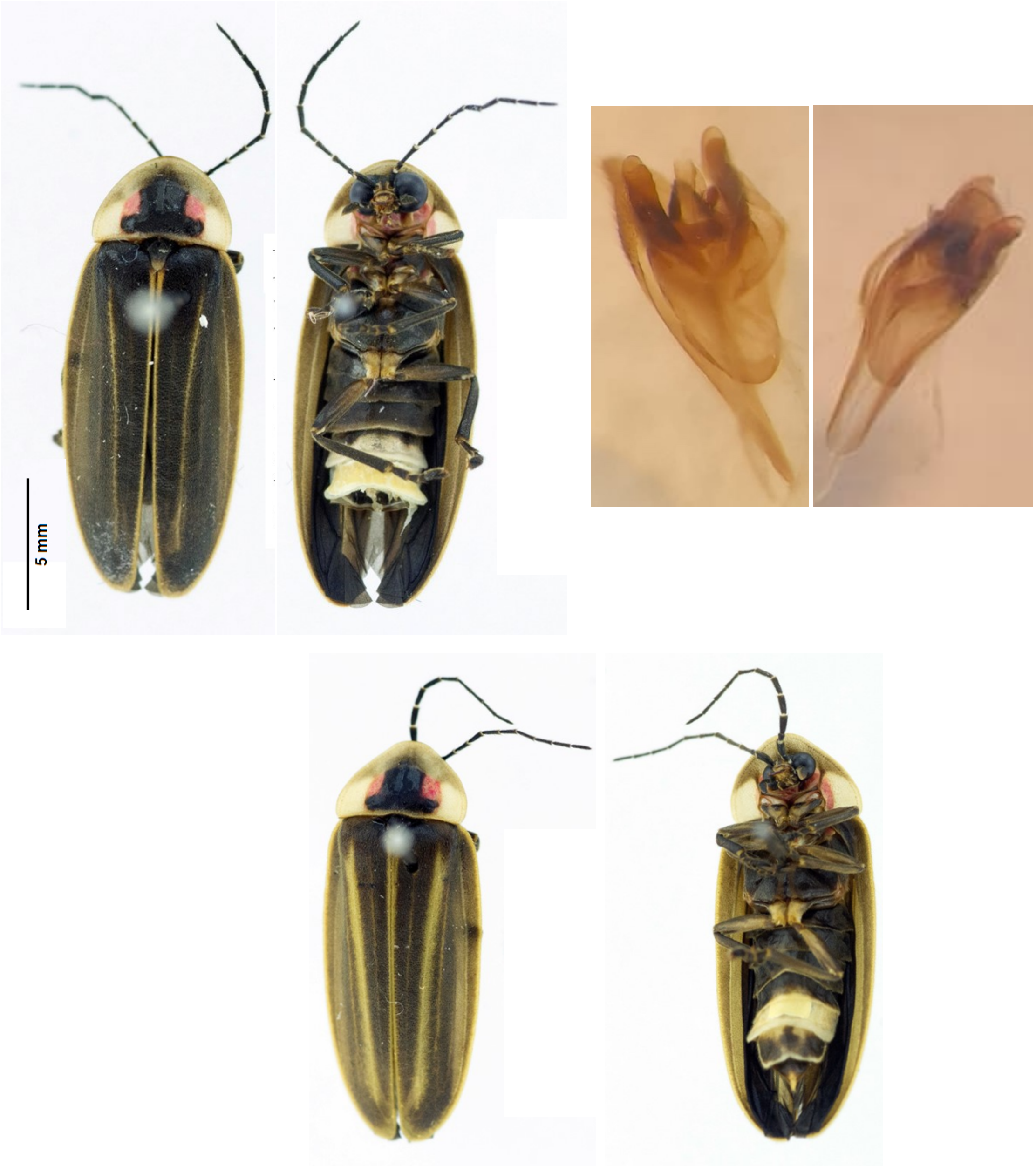


**Figure S5.** *Photinus congruus*. Upper panel: *male* dorsal and ventral view. Edeagos: dorsal, ventral and lateral views.  Lower panel: female dorsal and ventral view.

***Photinus sp7***

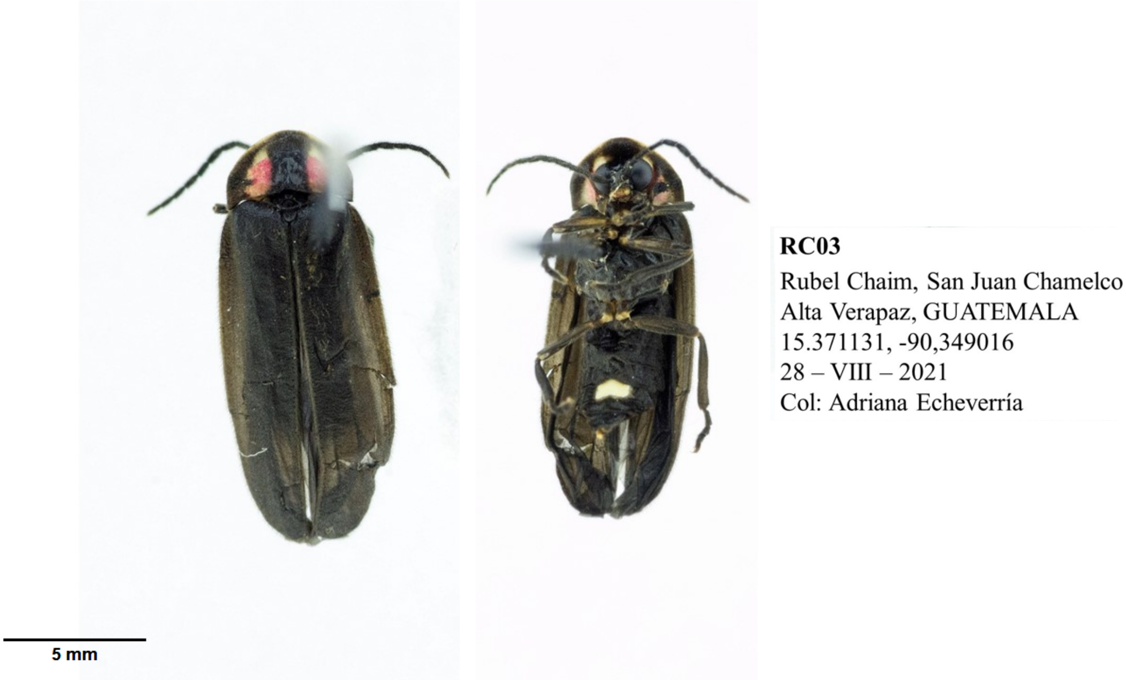


**Figure S6.** *Photinus sp7*. Female, dorsal and ventral view and collecting label.

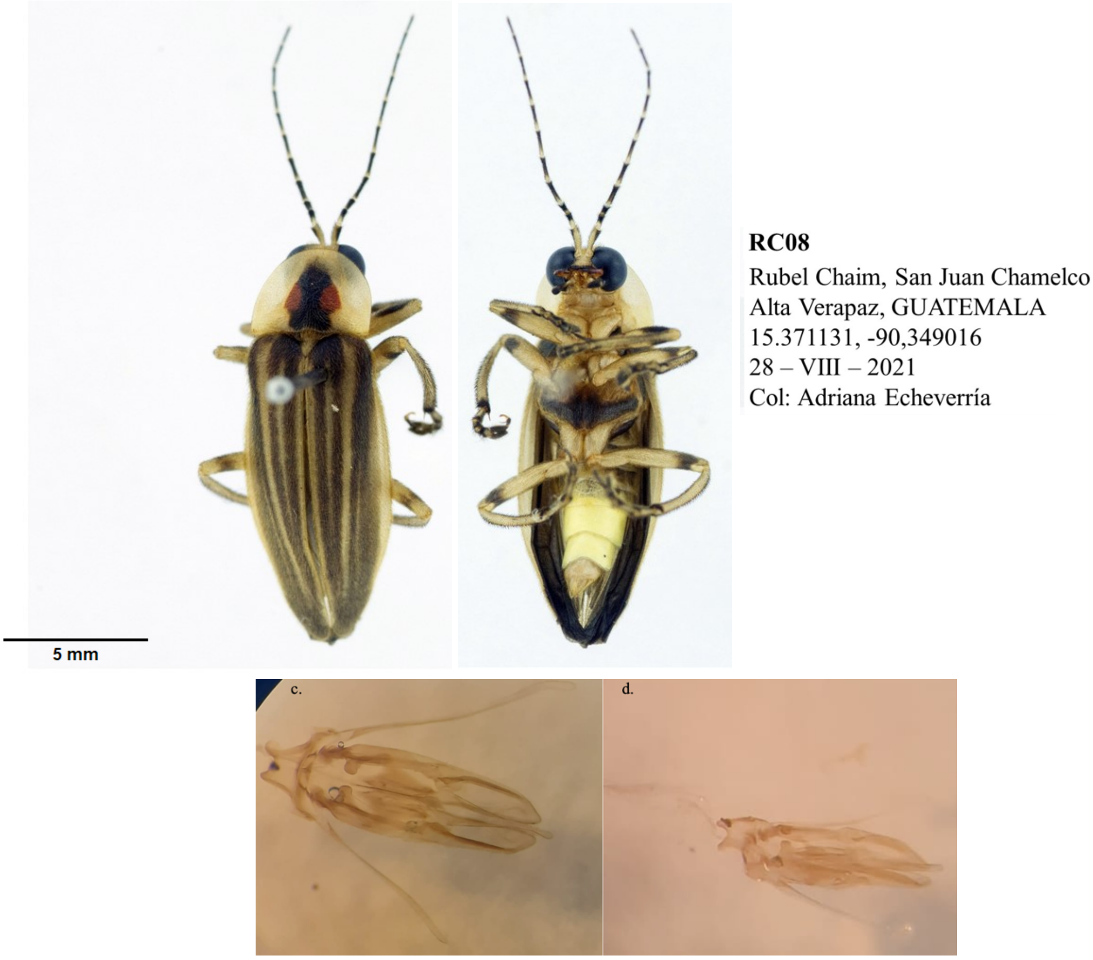

**Figure S7.** *Photuris sp1*. Upper panel: *male* dorsal and ventral view. Lower panel: Edeagos: dorsal and lateral views.

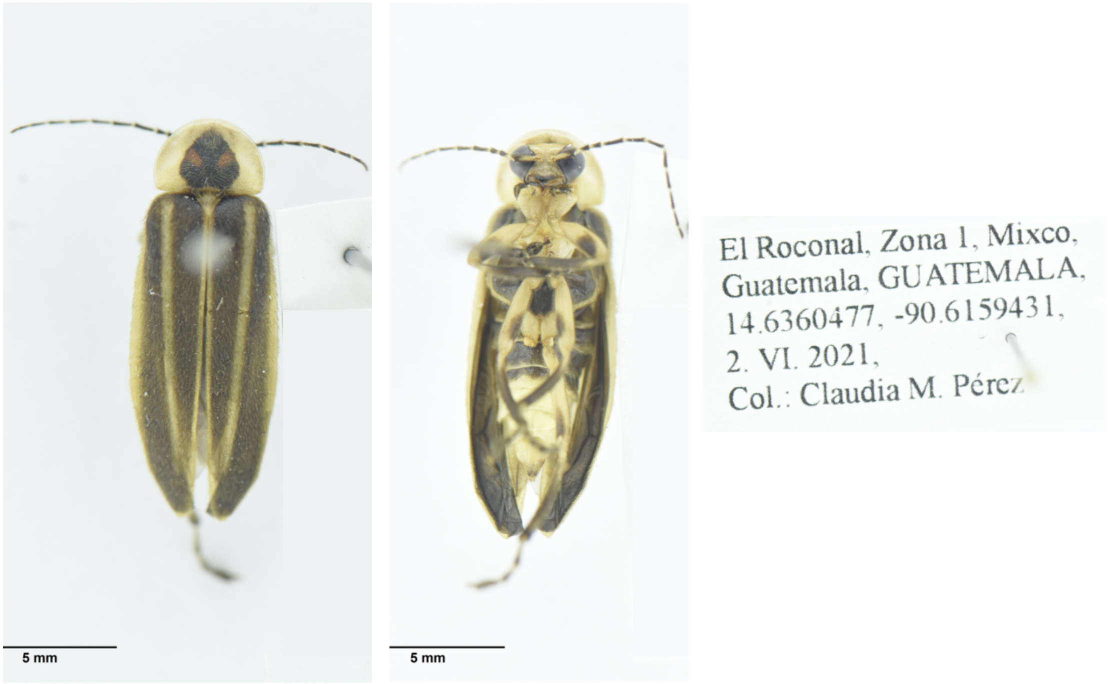


**Figure S8.** *Photuris sp2*. Male, dorsal and ventral view and collecting label.

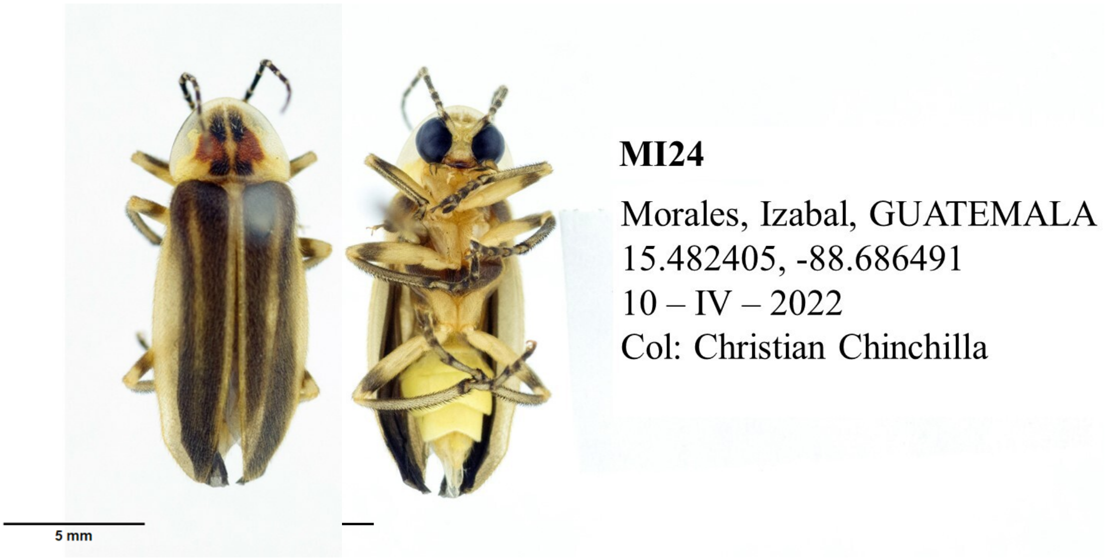


**Figure S9.** *Photuris sp3*. Male, dorsal and ventral view and collecting label.

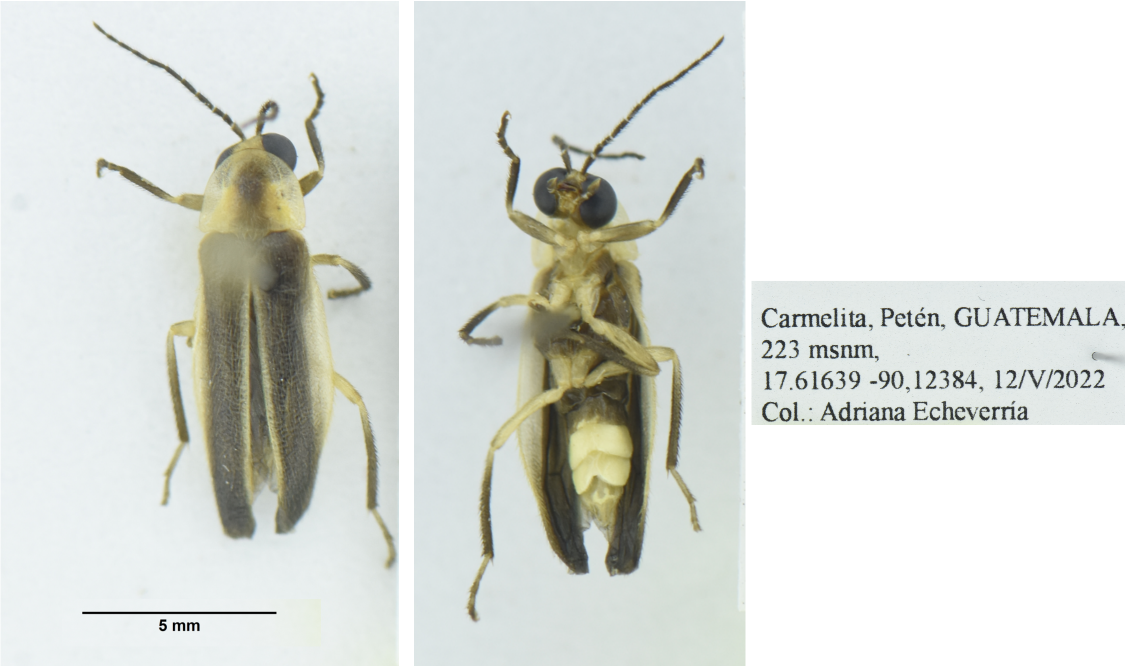


**Figure S10.** *Photuris sp4*. Male, dorsal and ventral view and collecting label.

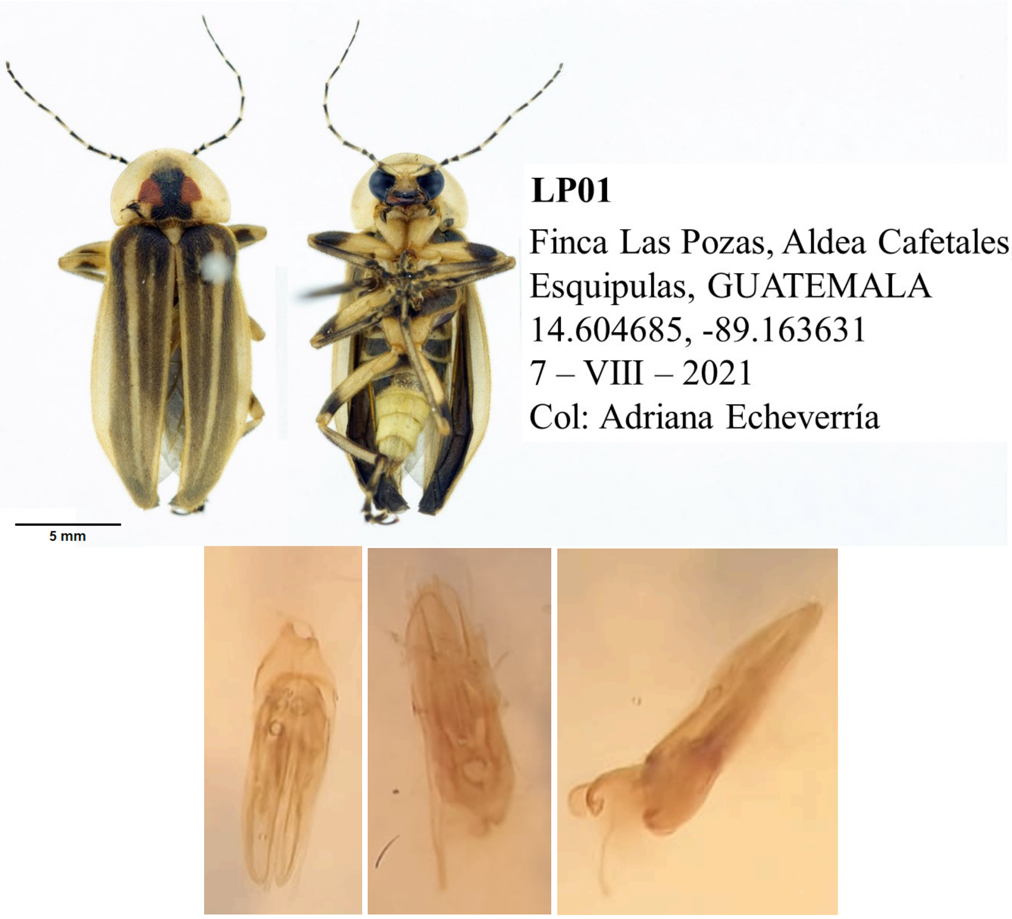


**Figure S11.** *Photuris sp5*. Upper panel: male dorsal and ventral view. Lower panel: Edeagos: dorsal, ventral and lateral views.

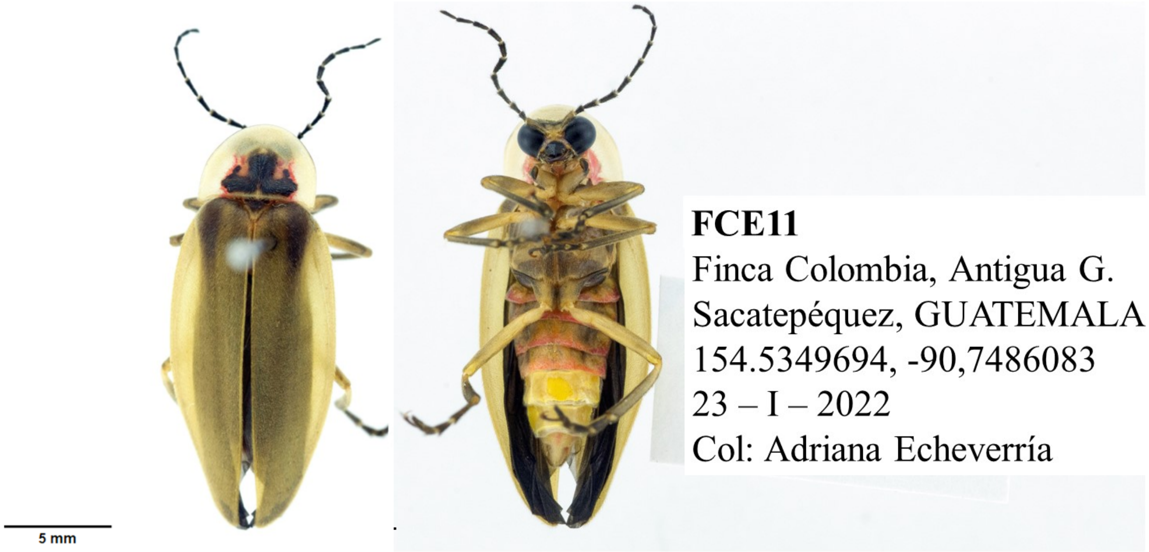


**Figure S12.** *Bichellonycha sp1*. Female, dorsal and ventral view including collecting label.

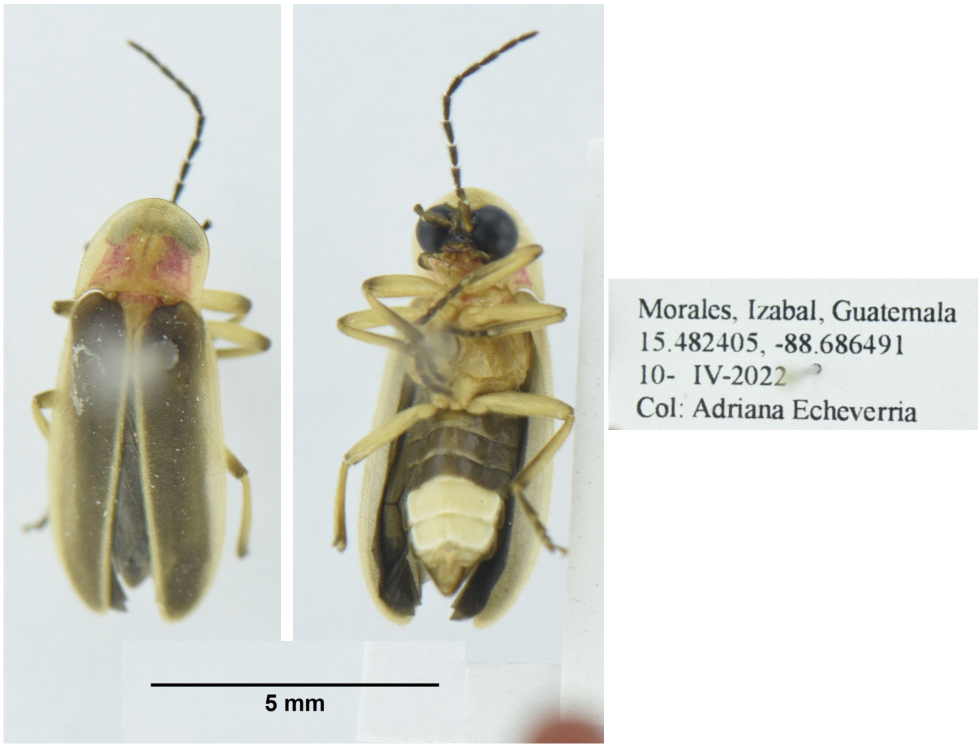


**Figure S13.** *Bichellonycha sp2*. Male, dorsal, ventral and collecting label.

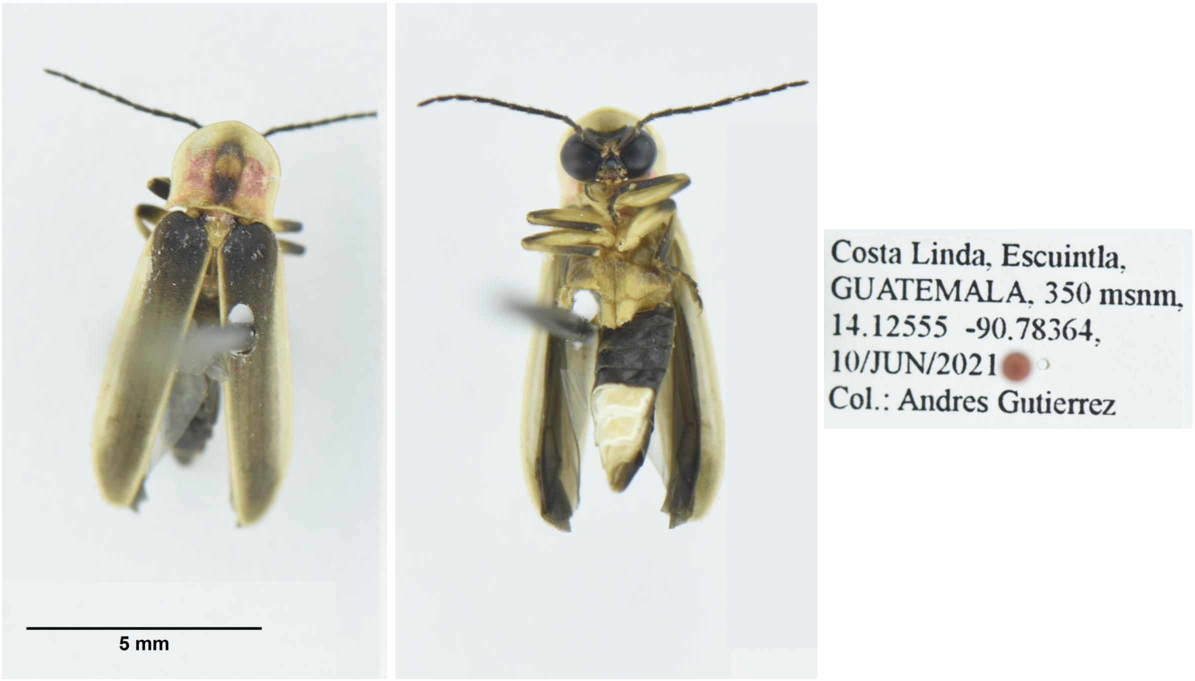


**Figure S14.** *Bichellonycha sp3*. Male, dorsal, ventral and collecting label.

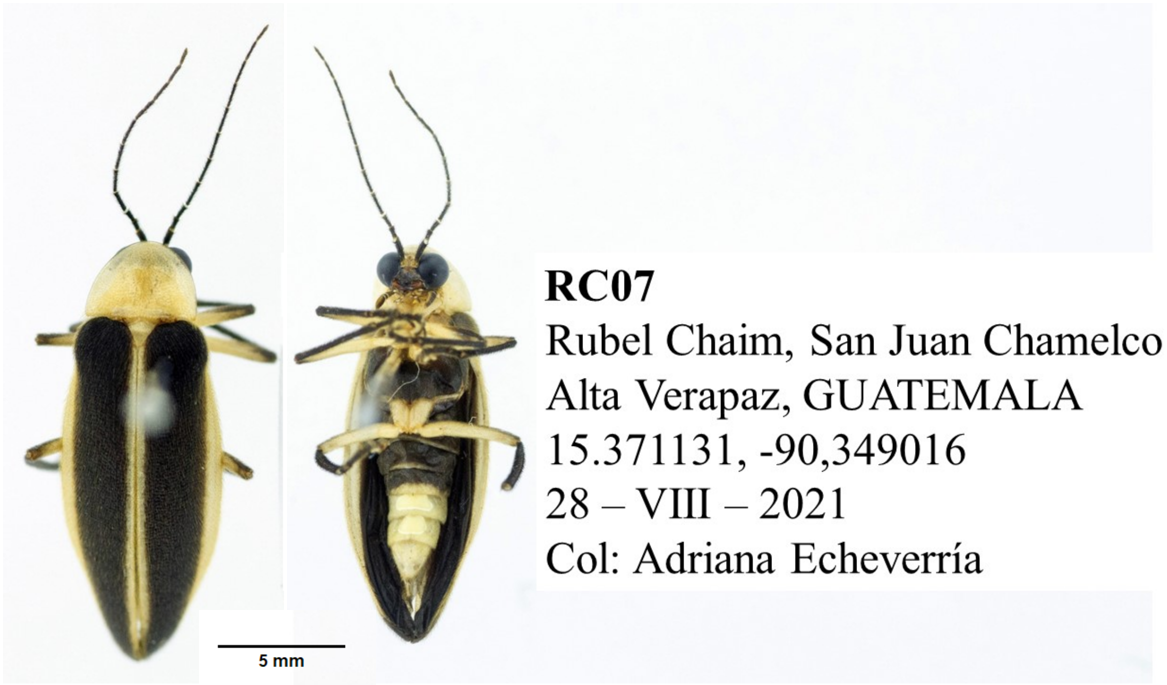


**Figure S15.** *Bichellonycha sp4*. Male, dorsal, ventral and collecting label.
